# Sensor localization using magnetic dipole-like coils: A method for highly accurate co-registration in on-scalp MEG

**DOI:** 10.1101/661678

**Authors:** Christoph Pfeiffer, Silvia Ruffieux, Lau M. Andersen, Alexei Kalabukhov, Dag Winkler, Robert Oostenveld, Daniel Lundqvist, Justin F. Schneiderman

## Abstract

Source modelling in magnetoencephalography (MEG) requires precise co-registration of the sensor array and the anatomical structure of the measured individual’s head. In conventional MEG, positions and orientations of the sensors relative to each other are fixed and known beforehand, requiring only localization of the head relative to the sensor array. Since the sensors in on-scalp MEG are positioned on the scalp, locations of the individual sensors depend on the subject’s head shape and size. The positions and orientations of on-scalp sensors must therefore be measured at every recording. This can be achieved by inverting conventional head localization, localizing the sensors relative to the head - rather than the other way around.

In this study we present a practical method for localizing sensors using magnetic dipole-like coils attached to the subject’s head. We implement and evaluate the method in a set of on-scalp MEG recordings using a 7-channel on-scalp MEG system based on high critical temperature superconducting quantum interference devices (high-*T*_*c*_ SQUIDs). The method provides accurate estimates of individual sensor positions and orientations with short averaging time (≤ 2 mm and < 3 degrees, respectively, with 1-second averaging), enabling continuous sensor localization. Calibrating and jointly localizing the sensor array can further improve the localization accuracy (< 1 mm and < 2.5 degrees, respectively, with 1-second coil recordings).

We demonstrate source localization of on-scalp recorded somatosensory evoked activity based on co-registration with our method. Equivalent current dipole fits of the evoked responses corresponded well (within 5.3 mm) with those based on a commercial, whole-head MEG system.

## 1. Introduction

On-scalp magnetoencephalography (MEG) has been shown in simulations to provide distinct advantages over traditional, low-*T*_*c*_ SQUID-based MEG. At closer proximity to the head –and thus to the neural sources– on-scalp MEG should be able to measure weaker signals as well as capture higher spatial frequencies compared to conventional MEG [1, 2]. In addition to smaller standoff, on-scalp MEG sensors - primarily optically pumped magnetometers (OPMs) and high-*T*_*c*_ SQUIDs - allow flexible sensing of the head; that is, the sensors can be moved (individually or in small units containing a few sensors) relative to each other in order for the sensor array to fit the head of individual subjects [3]. This is especially beneficial for studies on children, whose heads are significantly smaller than the one-size-fits-all helmets in most commercial MEG systems [4].

In general, translating MEG (sensor-level) signals to neural (source-level) activity requires co-registration of functional and structural data. An important step in this process is the reliable determination of the measurement/sensor locations relative to the subject’s head during the recording. In conventional MEG systems this is achieved by placing a set of small magnetic coils on the subject’s head and digitizing their positions with respect to landmarks (e.g., fiducials) on the head. Energizing the coils at different times and/or frequencies and detecting the distribution of the magnetic fields they generate (with the MEG system) allows accurate localization of the coils relative to the MEG sensor array [5, 6]. In order to localize the coils in such a way, the positions and orientations of the sensors relative to each other have to be known. This presents an issue when using flexible sensor arrays in on-scalp MEG. Because the sensors in such a system would be at least partially independently positioned, the sensors’ relative positions and orientations vary from subject to subject, and from session to session. Instead of a one-time calibration as used with rigid, whole-head sensor arrays, it is necessary to determine the sensor locations for each MEG recording session.

Measuring all the sensor positions and locations in a full-head array manually would be very time consuming and cumbersome, especially in arrays with high channel count. We have therefore developed and simulated the efficacy of a method for localizing independent MEG sensors with an array of small, magnetic dipole-like coils attached to the subject’s head [7]. Herein, we present the implementation of this sensor localization method in MEG recordings with a 7-channel high-*T*_*c*_ SQUID-based on-scalp MEG system. We furthermore validate its utility by using in source localization of somatosensory evoked fields.

## 2. Methods

### 2.1. Sensor localization

For an array of on-scalp MEG sensors recording a set of magnetic dipole-like coils (e.g., head position indicator, HPI, coils), the signal generated at the *k*th magnetometer by the *j*th magnetic dipole whose moment is 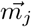 can be defined as

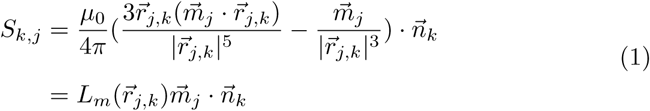

where *L*_*m*_ is the lead field, 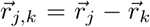 a vector defining the location of the dipole *j* relative to sensor 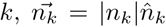 a vector combining the orientation 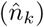 and sensitivity (|*n*_*k*_|) of sensor *k*, and 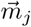 the magnetic moment of dipole *j*.

The position and orientation of a magnetic dipole is fit to recorded data 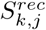 by finding the dipole location that minimizes the residual variance between the data and the calculated signals.

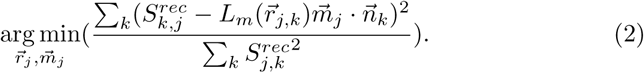

As described in [7], the standard coil localization procedure can be adapted to determine the position and orientation of an individual MEG sensor with respect to an array of coils by simply swapping the roles of magnetometers and dipoles:

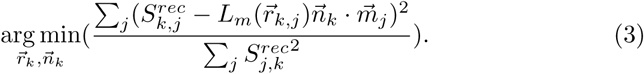

The on-scalp MEG system used here employs seven sensors that are fixed relative to each other in a single cryostat [8]. When multiple sensors are fixed relative to each other it is, in principle, possible to improve their localization by taking into account the array’s geometry [7]. Instead of solving eq. 3 for each sensor individually, the array can be combined into a single localization routine, wherein a single rigid transformation (rotation and translation) is applied to the whole sensor array. The number of parameters to be estimated is thus reduced by a factor of 7 compared to localizing the sensors individually. In this case, eq. 3 is replaced by:

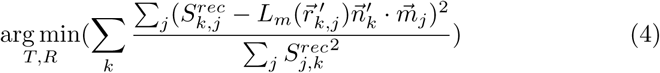

where *T* and *R* describe the 3-dimensional translation and rotation applied to the entire array, 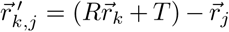 is the location of the rigidly transformed position of sensor *k* relative to dipole *j*, and 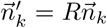 the rigidly transformed sensitivity vector.

To reduce the impact that noisy sensors can have on the localization accuracy, the sensors can be weighted according to their signal-to-noise ratio when summing the residual variances in eq. 4.

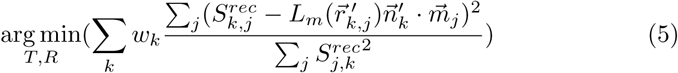

where 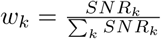 is the weight applied to the k-th sensor.

### 2.2. Measurement setup

The sensor localizations described here were performed as part of a set of MEG recordings at the National MEG Facility (NatMEG) at the Karolinska Institutet in Stockholm, Sweden. The main aim of the recordings was to compare and contrast recordings with a 7-channel high-*T*_*c*_ SQUID-based on-scalp system [8] to recordings with a commercial, whole-head system - in this case, a 306-channel Elekta TRIUX system (Elekta Neuromag Oy). Several different experimental paradigms were recorded in five neurotypical subjects (4 male and 1 female, ages 30-49). For each session the same paradigm was first recorded on a subject with the commercial MEG system, followed by the on-scalp MEG recording. All experiments were approved by the Swedish Ethical Review Authority (EPN 2018-571-31-1) and conducted in compliance with national legislation and the code of ethical principles defined in the Declaration of Helsinki. All participants gave informed consent.

Ten dipole-like head position indicator (HPI) coils of the TRIUX system were used both in the head localization as part of the conventional MEG recordings and in the sensor localization as part of the on-scalp recordings. The coils were driven at frequencies from 537 to 987 in steps of 50 Hz. The frequencies were chosen relatively high in order to spectrally separate them from neural activity (including high frequency components up to 500 Hz). The frequency steps are chosen such that potential intermittent-frequency artefacts would coincide with the power line harmonics (50 Hz in Sweden), which are filtered as part of the standard preprocessing and therefore do not require any additional treatment.

The recordings were divided into blocks of stimulations with the coils energized for 10 to 30 seconds before and after each block. This was done as a cautionary measure to prevent potential artifacts from the coils to corrupt the MEG recordings. Recording before and after each stimulation block also allowed monitoring if/how the head moved.

The subjects were recorded seated with their heads comfortably stabilized using vacuum pillows (without being completely immobilized). To further minimize head movements during the coil recordings, the subjects were instructed to keep their head still. For each paradigm (in some cases two paradigms with similar neural activation) a coarse region of interest was determined prior to the recording session based on knowledge about the expected activity and/or previous recordings on the same subject using the same or a similar paradigm. The coils were then distributed closely around the region of interest, while maintaining sufficient room for placement of the cryostat. In order to minimize relative movements between coils, nine coils were fixed to small plastic plates (three coils per plate) that were roughly shaped to fit to the subject’s head. The tenth coil was then fixed to the head individually. Figure 1 shows a set of coils arranged around a region of interest on an EEG cap on one of the subject’s head. The red tags mark the different target locations for the on-scalp system. The coils, head shape and target location tags were digitized using a AC electromagnetic tracking system Polhemus Fastrak (Polhemus, Colchester, VT 05446, USA).

**Figure 1:**
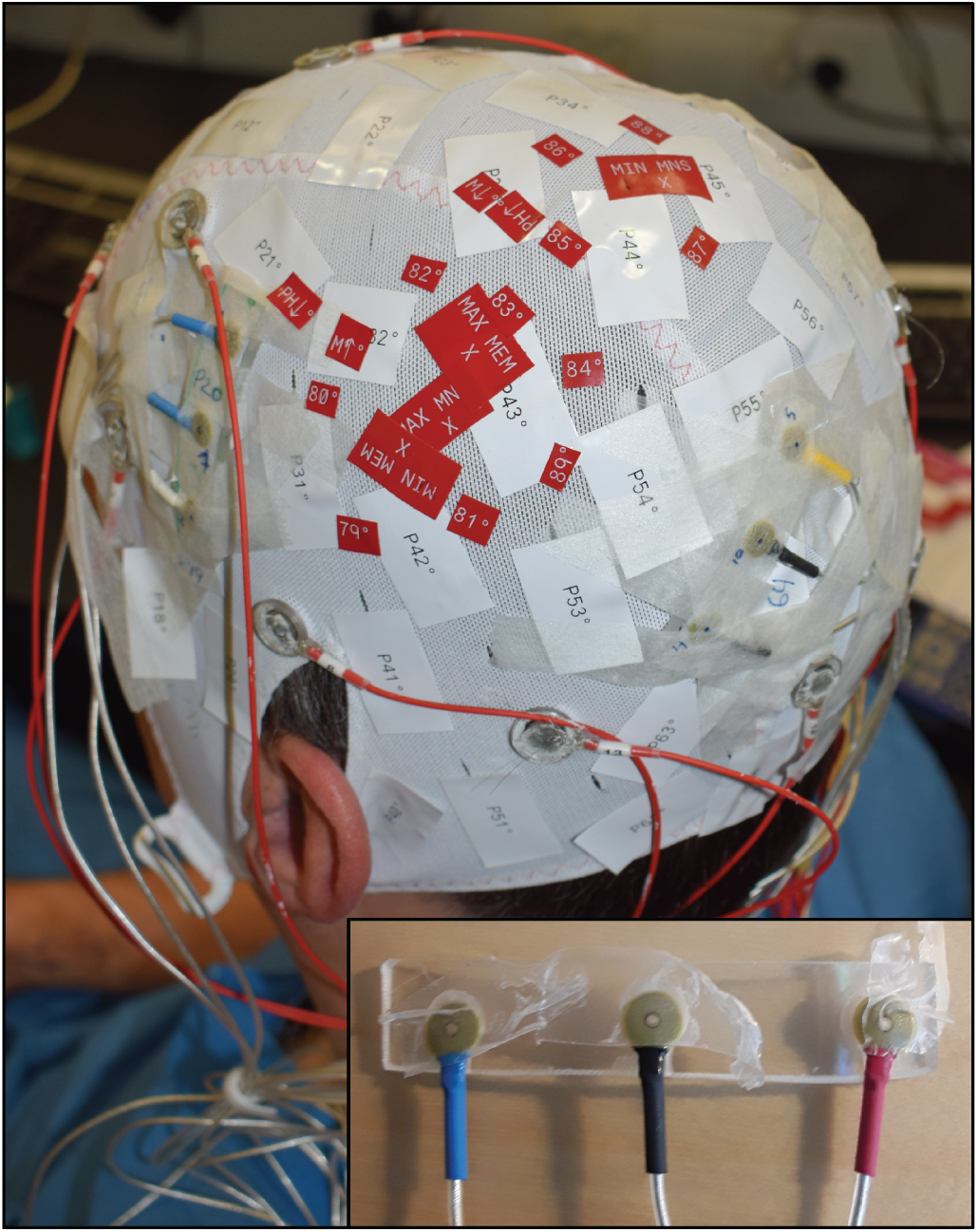
Photograph showing HPI coils attached to a subjects head. Three triplets of coils (each attached to a rectangular plastic holder) can be seen surrounding a region of interest marked by red tags that indicate measurement locations. Inset: a plastic holder with three HPI coils attached.

At the beginning of each recording the subject was recorded in the TRIUX system. These recordings were used to localize the underlying neural activity and project the resulting neuromagnetic fields onto the scalp surface. Such field maps were used to guide the placement of the cryostat (i.e., the red markers in Fig. 1) for each experimental paradigm and subject [9, 10]. More importantly for localizing the sensors, the whole-head recordings were used to determine the positions, orientations, and magnetic moments of the coils relative to each other and to the head via traditional head localization [5]. HPI coil locations and orientations obtained thus were used for the ensuing on-scalp recordings. Only coil locations where the goodness of fit exceeded 0.98 were used in the sensor localization.

The sensor fits were performed in MATLAB R2015a (Mathworks, Natick, MA, USA) using the FieldTrip toolbox [11]. The coil amplitudes were extracted from the data via multitaper frequency transform using Slepian tapers and used in a linear grid search to provide a starting point for the non-linear fit. Finally, the sensor locations were fitted to the extracted coil amplitudes by solving eq. 3 using unconstrained optimization (quasi-newton algorithm) with the starting point obtained from the grid search.

When fitting the sensors jointly, the known layout of the sensor array is rigidly aligned to the individually fitted sensor locations using an iterative closest points (ICP) algorithm that was modified to minimize distances between corresponding point pairs (that is, points corresponding to the same sensor) rather than closest points. The resulting transformed sensor array then serves as starting point for a non-linear fit.

### 2.3. Evaluation

Defining the performance of the sensor localization is not straightforward in a realistic measurement setup, like the one we present here, wherein the “ground truth” (i.e., the true sensor locations relative to the head) is not known with arbitrary precision. Generally, the accuracy of the fitted locations are affected by a combination of random errors (e.g., due to sensor noise), systematic errors (resulting from, e.g., errors in the coil positions) and variations in the true location (resulting from head movements).

#### 2.3.1. Random errors

Assuming head movements are negligible during a single (30 second) recording, we estimate the effects of random errors. We split each 30-second coil recording into multiple shorter segments, each of which was independently used to localize the sensors. Variations in an individual sensor’s location over segments were then used to provide an estimate of the sensor localization accuracy. To this end, we define 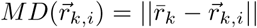 as the euclidean distance of the i-th segment’s fitted position 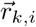 from the mean location 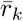 over all such segments. Describing the spread of the sensor locations around the mean *MD* provides an estimate of random errors - and thus the location accuracy. Similarly, we define 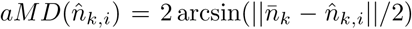 as an estimate of the angular accuracy (i.e., the segment-by-segment angular deviation of the corresponding sensor orientations from the mean orientation over segments 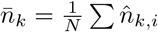).

#### 2.3.2. Systematic errors

One limitation to these metrics is that they do not provide information about systematic errors that would result in a shift in the mean position. Furthermore, despite subjects’ efforts to minimize head movement during coil recordings, the possibility of small movements cannot be excluded - the subjects heads were comfortably stabilized with vacuum pillows, but not immobilized. These issues can be dealt with by taking advantage of the fact that the sensors are housed in a common cryostat, i.e., fixed relative to each other. The distances between the (true) sensor locations are thus constant and independent of head movements. Localization errors can therefore also be estimated by comparing the distances between the fitted sensor locations with those from the known layout of the sensor array. To this end, we estimate a relative localization accuracy as the average deviation of the distances between the estimated sensor locations from the distances derived from the known layout:

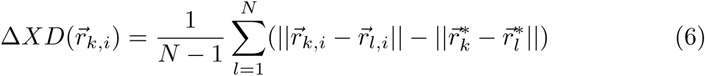

where 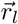 and 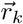 denote the positions of the localized sensors l and k, 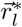 and 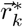 their respective positions according to the reference (e.g., the system design), and N=7 the number of sensors. The sum is divided by N-1 because the term for l=k is always zero. This metric is only useful for evaluating individual sensor fits because distances between sensors are constant and determined by the sensor array when jointly localizing the sensors (because the positions are a result of rigidly rotating and translating the sensor array). Analogously, we can estimate the relative localization accuracy with respect to the orientation as the average deviation of the angles between the estimated sensor orientations from the angles between the reference sensor orientations:

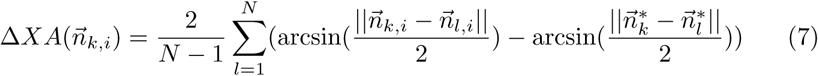

where 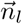 and 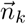 denote the orientations of the localized sensors l and k and 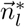 and 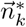 their orientations according to the reference (e.g., the system design).

#### 2.3.3. Head movements

Localizing sensors from shorter coil recordings/segments is favourable when trying to detect - and compensate for - head movements as it enables estimation of recording positions with higher temporal resolution. This is how head movements are conventionally detected/tracked: the sensor locations with respect to the head are estimated at multiple time instances and compared to the initial position. In order for us to investigate how the accuracy of the sensor localization depends on the time the coil signals are recorded, *MD, aMD* and Δ*XD* were computed for different segment lengths *t*_*trial*_ between 1 and 10 seconds. For each segment length, the 30 seconds coil recording was split into n = 30/*t*_*trial*_ consecutive trials.

#### 2.3.4. Source localization

Finally, we tested the usefulness of our sensor localization procedure in localizing neural activity. The MEG experiments included recordings of somatosensory evoked fields (SEFs). Using our sensor localization method for co-registration of the on-scalp data, source localization of the N20m-component was performed and compared to source localization using the conventional MEG data recorded with the TRIUX system. Because of the small coverage of the on-scalp system we recorded at four separate locations (aimed to capture the dipolar field pattern of the N20m-component) and combined the resulting data. One sensor was excluded due to excessive noise, resulting in 24 individual sensor locations. The same experimental paradigm - electric stimulation (below motor threshold) of the median nerve with 360 ms inter-stimulus interval and 1 000 repetitions - as well as preprocessing - bandpass filter between 5 and 200 Hz with bandstop filters applied at 50 Hz and harmonics, 50 ms pre- to 200 ms post-stimulus epochs, baseline correction (−50 to 0 ms baseline window), and time-locked averaging - was used for the recordings with both systems. For comparability, only the magnetometers were used for the dipole fit with the TRIUX system.

## 3. Results

The Fourier spectrum of a coil recording is shown in Fig. 2. Clear peaks with a signal-to-noise ratio (SNR) on the order of ∼ 10^2^ are visible at the coil frequencies. An example of a sensor localization based on an 10-second trial can be seen in Fig. 3. In this case, the fitted sensor positions and orientations match well with the design of the sensor array (all pairs being within 0.5 mm and 2 degrees of the design) [8].

**Figure 2:**
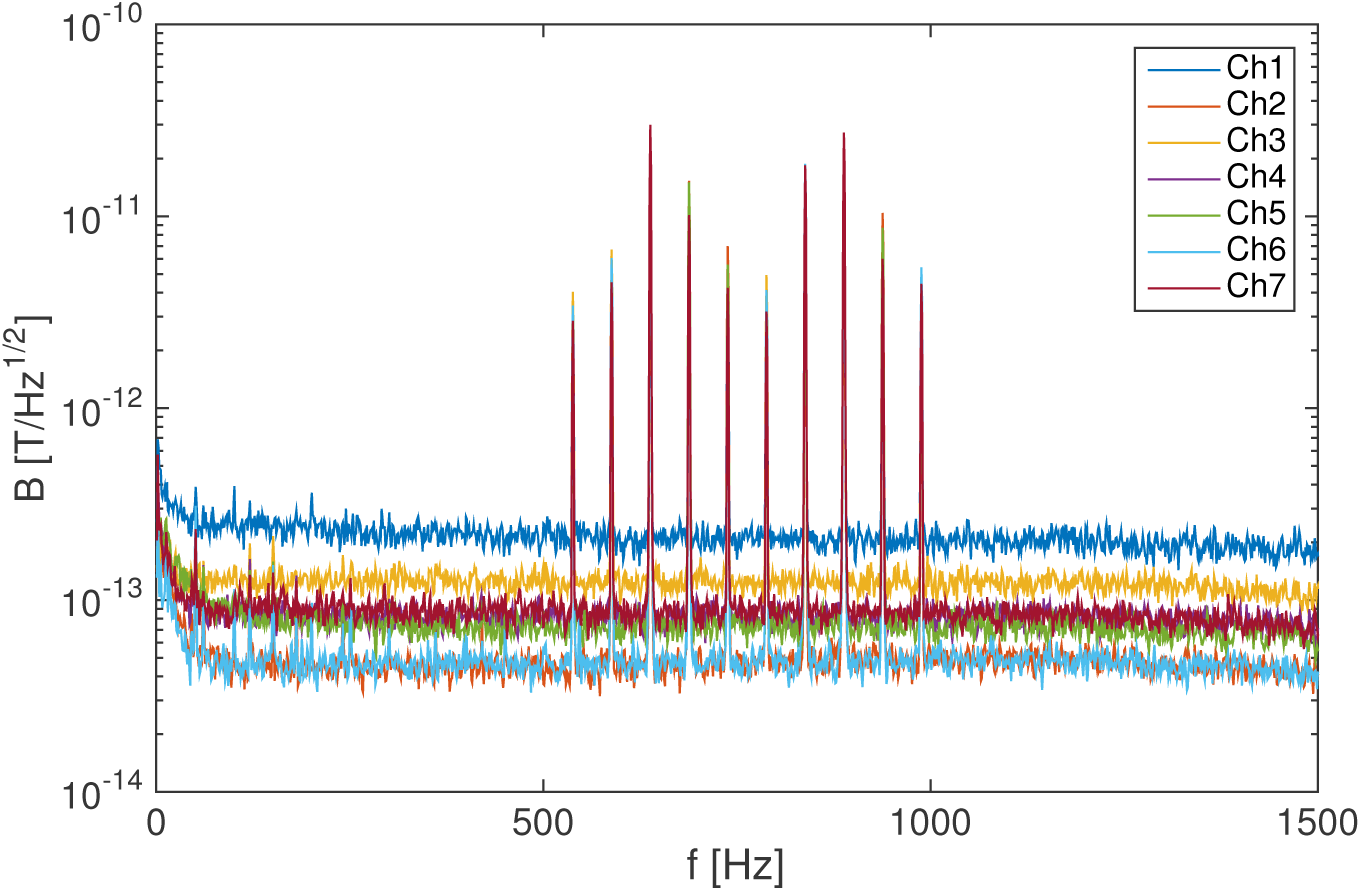
Spectrum of the measured magnetic fields showing peaks at the coil signal frequencies.

**Figure 3:**
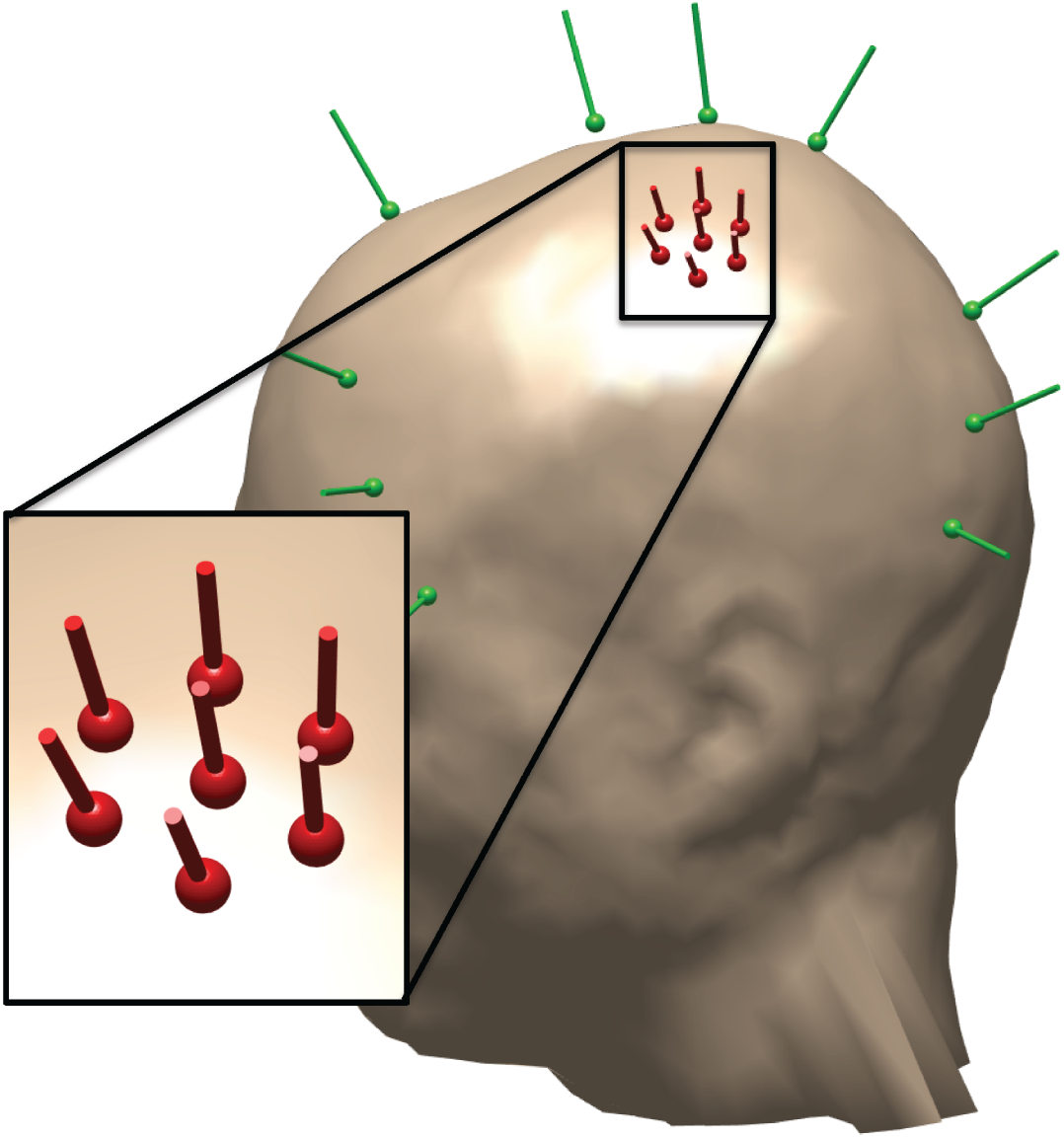
Example of individually fitted sensor locations and orientations (red). Magnetic dipoles from the coils are shown in green.

In some recordings, individual sensors trapped flux, which led to a strong increase in noise (∼10× higher white noise and a shift in the 1/f-like noise knee from 10 to 500-1000 Hz). Localization of these noisy sensors was severely degraded - with errors on the order of centimeters. However, with such high noise data from these sensors was not useful for the MEG recordings and the sensor localization therefore inconsequential.

Fig. 4-a shows the mean euclidean distances of the fitted sensor locations from the mean locations 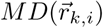 as a function of the duration of the coil recording segments *t*_*trial*_ used for the localizations. As expected, a clear correlation between the localization accuracy and the length of the coil recordings can be observed. With the exception of channel 1 (which exhibited high noise in the recording) all channels reach MD < 1 mm even with just 1-second recordings of the coil signals (channel 1 with four seconds or more). The mean angular deviations from the mean fitted sensor orientations 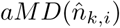- seen in Fig. 4-b - show a similar trend versus coil recording time. The orientation fits deviate from the mean by less than 3 degrees with one second of coil signal recording.

**Figure 4:**
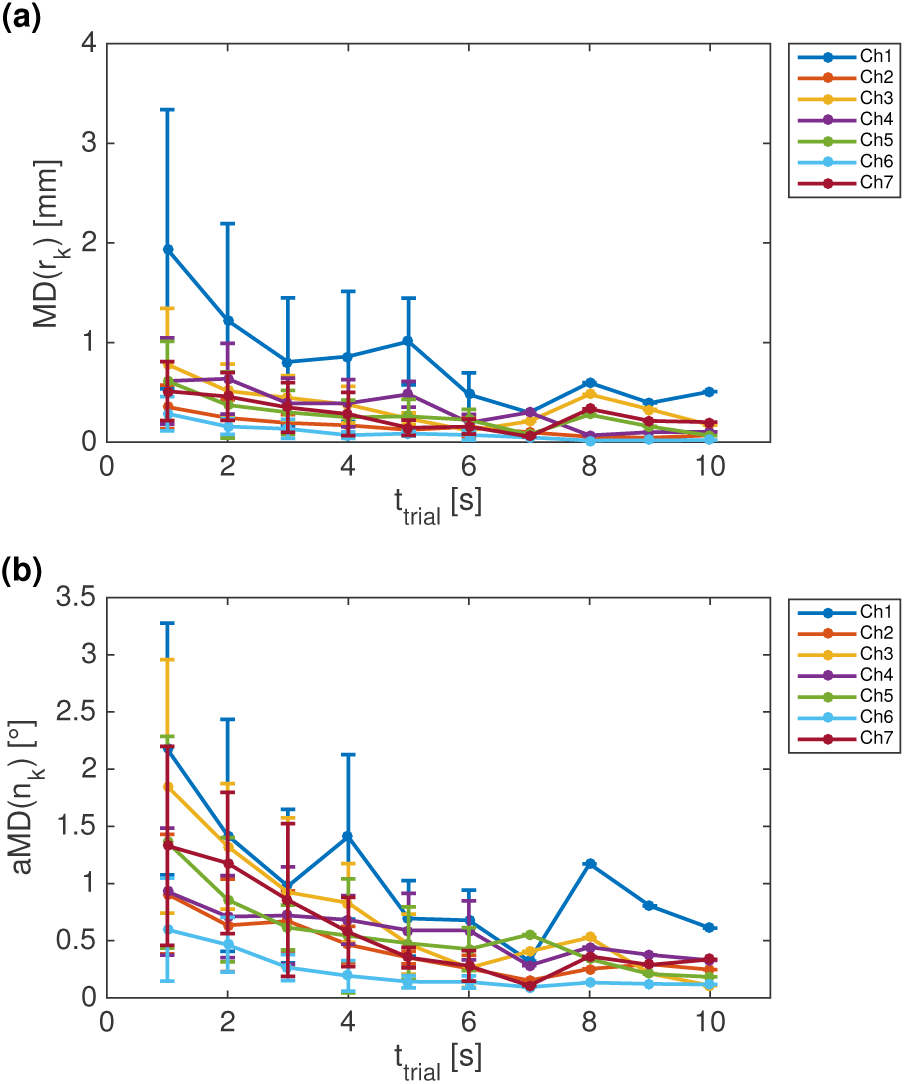
Sensor localization accuracy. a) Mean distance from the mean location 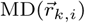 as a function of the segment length. b) Mean angular deviation from the mean orientation 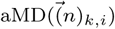 for different segment lengths. Error bars indicate one standard deviation.

Fig. 5-a shows the mean differences of the distances between the fitted sensors from the distances between sensors in a reference array, 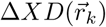. In this case, we used the design of the system as the reference and again present results for different lengths of coil recording segments *t*_*trial*_. On average all channels differ by less than 1 mm from the design already with 1-second coil recordings. With increasing *t*_*trial*_, the mean 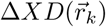 converge to values < *±*0.4 mm. These can be assumed to stem from a combination of systematic errors and small deviations between the actual sensor array and the design. As before, the decrease of the standard deviation (i.e., the segment-by-segment spread) with longer coil recording time indicates a decrease in random localization errors. The mean differences of the angles between the fitted sensors from the angles between the sensors in the design of the system 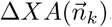, seen in Fig. 5-a, show a similar decrease in standard deviation with increasing coil recording time. With 1 second coil recordings all channels differ by ∼2 degrees or less from the design of the system.

**Figure 5:**
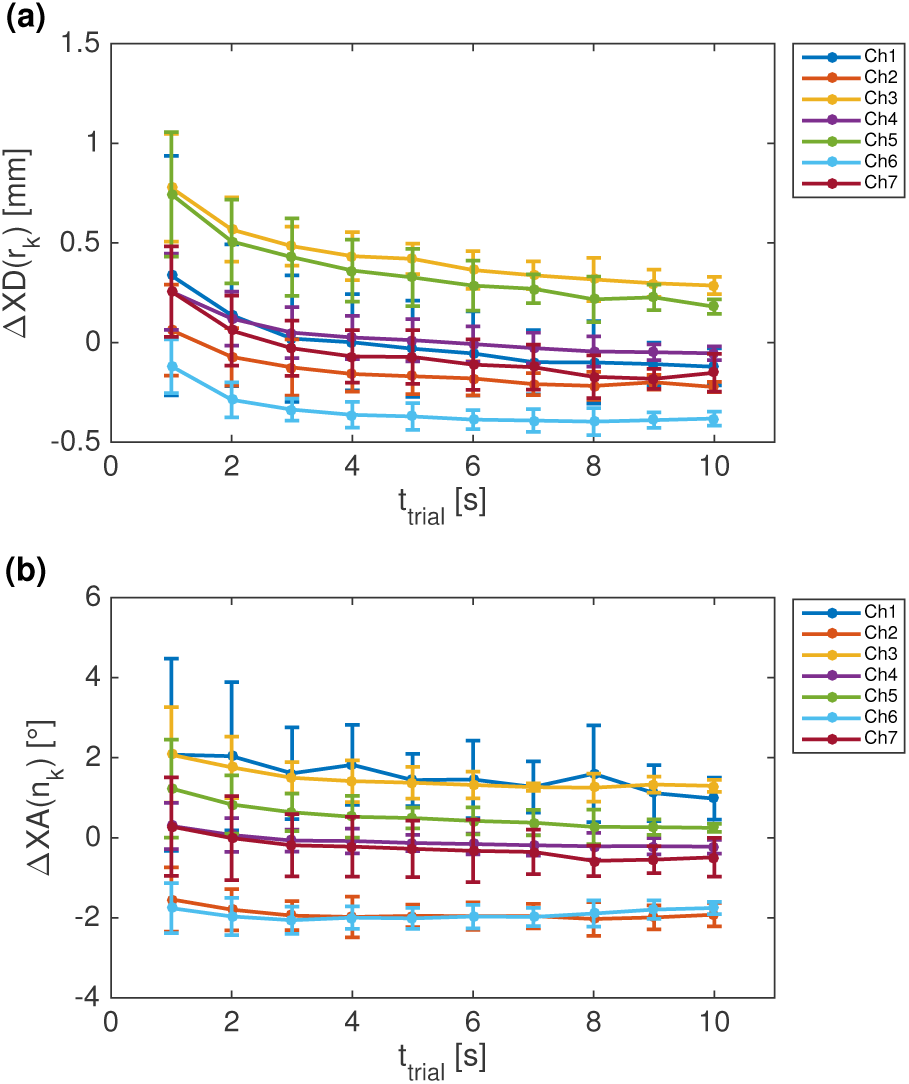
Pairwise sensor localization accuracy, with the cryostat design as the reference. a) Mean difference in distance to the other sensors 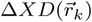. b) Mean difference in angle to the other sensors 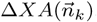.

Using short segments, it is possible to continuously monitor the sensor locations in order to detect movements of the head relative to the sensors. Head movements manifest themselves as a shift and/or rotation of the whole sensor array between segments. An example of a head movement captured with 2-second coil recordings can be seen in figure 6. In this case, the subject’s head moved approximately 2 mm upwards during a stimulus session.

**Figure 6:**
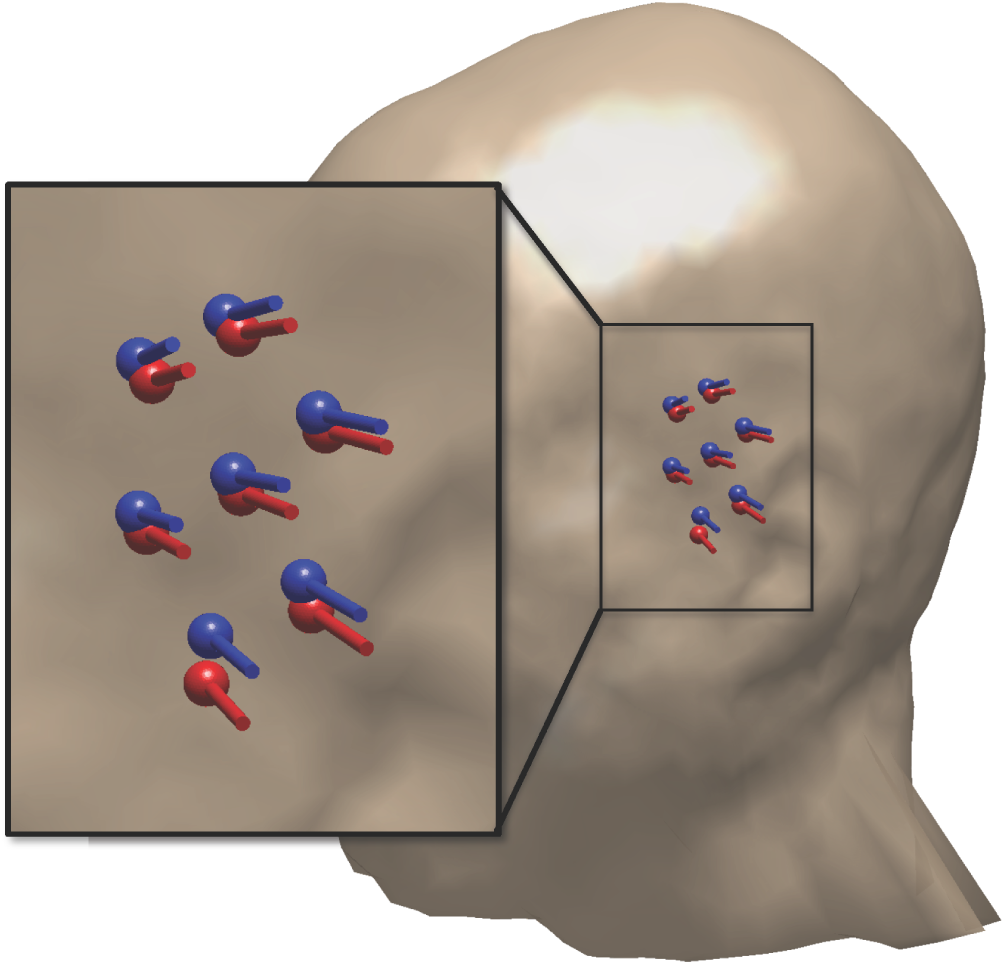
Successive sensor localizations (red and blue) showing head movement (∼ 2 mm) between coil recordings.

Distances from the mean location 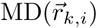 as well as angular deviations from the mean orientation 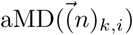 when localizing the sensors jointly are shown in Fig. 7. The joint localizations were performed on the same data used to individually localize the sensors in Fig. 4. Both 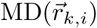 and 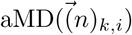 show a similar trend as when localizing the sensors individually. Compared to the individual localization, the noisier sensors show significant improvement (especially in 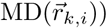 while the lower noise sensors worsen. However, the spread in location and orientation around the mean decreases in general, indicating an overall improvement in localization accuracy. This is especially pronounced in case of the location: with one second of data, all sensors exhibit 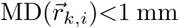 and 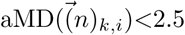 degrees (compared to ≤ 2 mm and <3 degrees, respectively, when localizing them individually). The joint localizations shown here were performed using the sensor positions obtained via individually localizing the sensors with a 10-second coil recording to define the sensor array.

**Figure 7:**
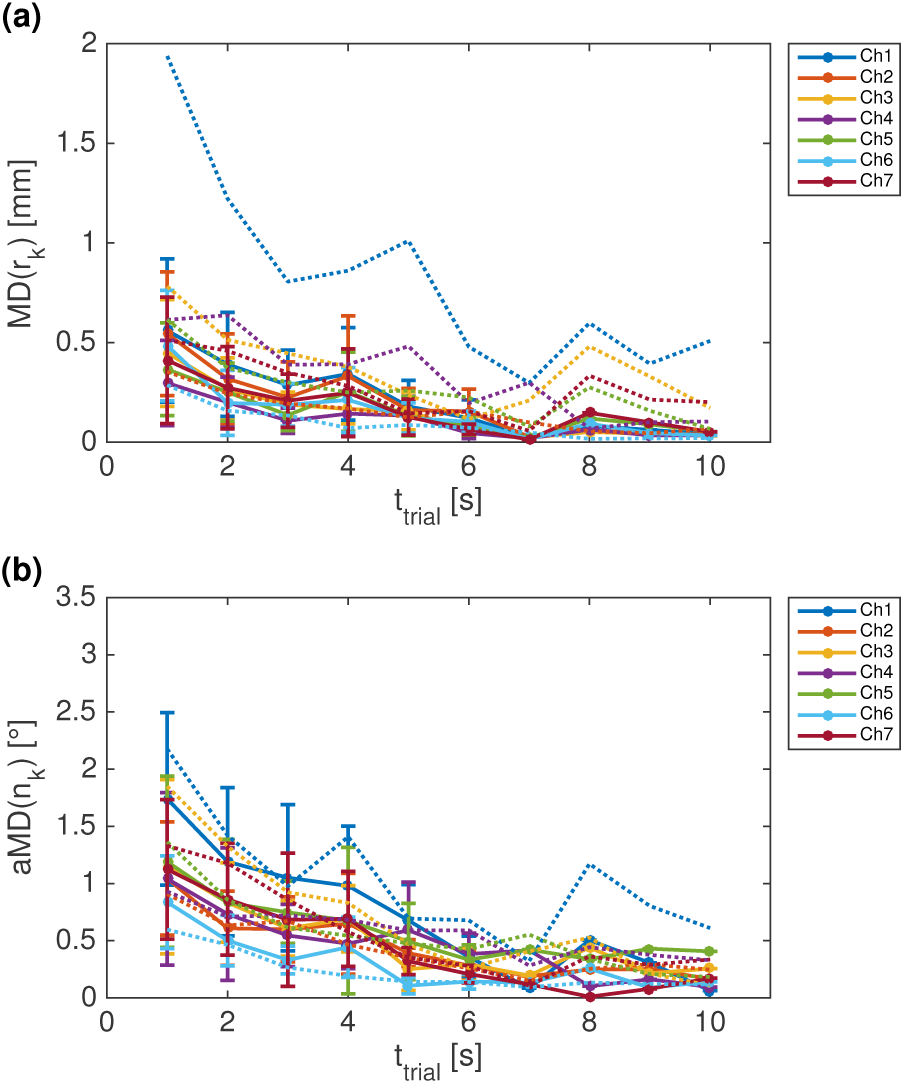
Joint sensor localization accuracy using the sensor locations obtained from 10-second coil recording individual localization as rigid sensor array. a) Mean distance from the mean location 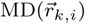 as a function of the segment length. b) Mean angular deviation from the mean orientation 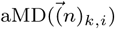 for different segment lengths. Error bars indicate one standard deviation. For reference, we include the mean of the corresponding deviations that were obtained when localizing the sensors individually as dotted lines.

Weighting the sensors according to SNR to reduce the impact of noisy sensors (here, e.g., Ch1) did not result in an improvement in accuracy. In fact, the average accuracy for long coil recordings when localizing SNR-weighted sensors was worse compared to localizing equally weighted sensors.

Dipole fits of the N20m-component recorded on-scalp and conventionally can be seen in Fig. 8. The two dipoles are 5.3 mm apart, which is within the localization accuracy of conventional whole-head MEG systems [12, 13].

**Figure 8:**
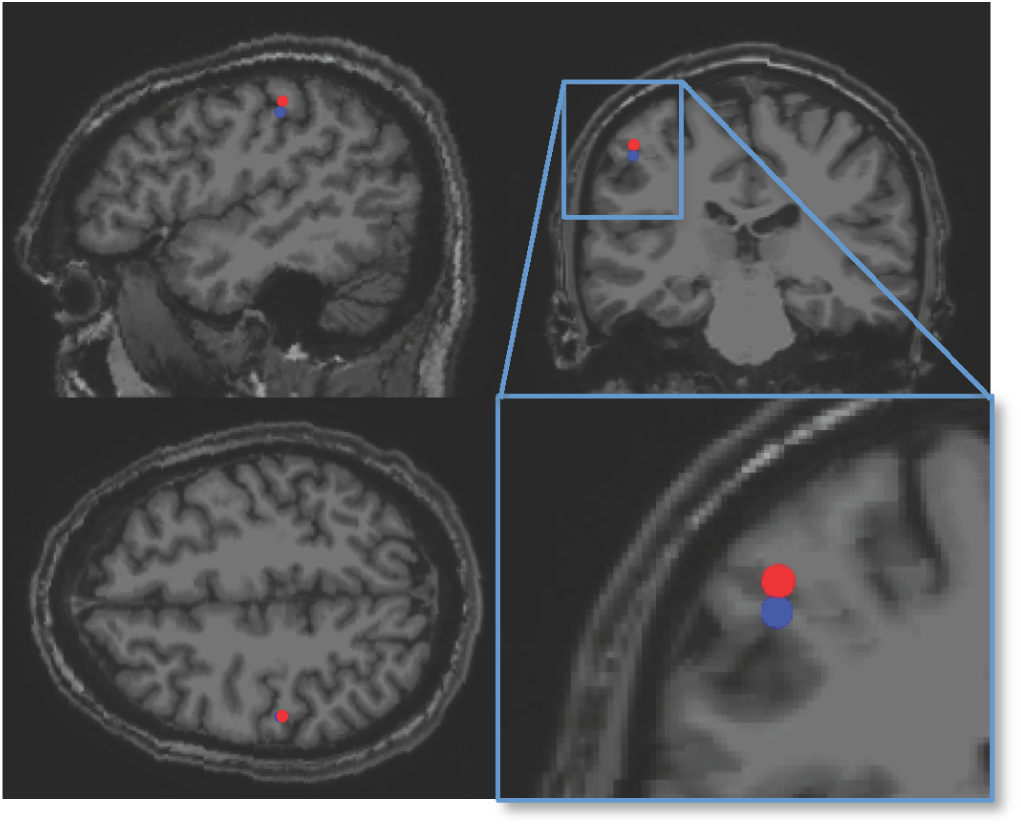
Dipole fits of N20m component based on on-scalp (red) and conventional (blue) MEG recording. The on-scalp dipole fit was performed using individually localized sensor positions estimated with our method.

## 4. Discussion

With ≤ 2 mm and < 3 degrees for 1-second coil recordings, the sensor localization method described here reaches significantly higher accuracy than what has been suggested as required for on-scalp MEG (<4 mm and <10 degrees, according to [14]).

An advantage of our method is that it allows for continuous co-registration in parallel with the MEG recording. Movements of the subject’s head during the MEG recording can thus be detected and accounted for, similarly to continuous head localization used in commercial whole-head MEG systems. The measurements shown here were a first practical attempt of using the method described theoretically in [7]. However, as the experimental session was performed in parallel with other on-scalp MEG experiments, we erred on the side of caution by turning the coils off during stimulations (in order to avoid the possibility that they would generate artifacts that might compromise the MEG recordings). While it remains to be experimentally verified, it is likely that our method can be used during a stimulus or other experimental protocol because the coil recordings showed no interference at frequencies below 500 Hz (see Fig. 2). Furthermore, in cases where neural signals of interest coincide with the coil frequencies, it is trivial to change the coil frequencies to avoid potential interference (if the neural frequencies of interest are known). The upper limit for the coil frequencies is strictly set by the Nyquist frequency (half of the sampling frequency, in this case 5 kHz/2 = 2.5 kHz) and generally should be kept well below any low-pass filters used by the data acquisition system (e.g., anti-aliasing filters, in our case 1 600 Hz).

Taking advantage of the fixed geometry of the sensor array to jointly localize the sensors proved useful. The increased accuracy at shorter segment lengths is especially important for continuous sensor localization. Furthermore, by using individually localized sensor positions from a longer coil recording to define the array geometry, the method is not limited to systems where the sensor array is rigid. For systems consisting of multiple individually positionable sensors [15, 16, 17] or units containing a few sensors [18], one can calibrate the sensor array at the start of a recording by carefully recording the coil signals for a longer duration of time (while minimizing head movement) and localizing the sensors individually. The calibrated array can then be used for fast, joint sensor localization. This, of course, assumes that the sensors are fixed with respect to one another for the duration of the recording.

Localized sensor positions and orientations were used to fit an equivalent current dipole to somatosenory evoked activity recorded sequentially at multiple locations. The estimated dipole position from the on-scalp recording was ∼4 mm from that which was estimated from the conventional MEG recording. This lies well within the 8-11 mm variability seen between different commercial, whole-head MEG systems [13]. Considering the differences in sampling between on-scalp and conventional MEG, it is also possible that the on-scalp system is differently sensitive to neural activity, as compared conventional MEG. Previous works by our group with a high-*T*_c_ SQUID [10] as well as by Zetter et al. [14] with OPMs also report differences between the N20m-components detected with on-scalp and conventional MEG systems.

The measurements reported here were part of a series of benchmarking recordings to compare an on-scalp MEG system [8] to a commercial, whole-head MEG system. It was therefore possible to use full-head recordings of the coil array on the subject’s head in order to reliably estimate the positions and orientations of the dipolar coils. This is, however, not a viable solution for on-scalp systems in general. The coil orientations should instead be inferred from other measurements. Flat coils with markers to digitize the orientation as part of the head-digitization would be able to solve this issue in the future.

## 5. Conclusion

We have presented a method for localizing MEG sensors with the help of magnetic dipole-like coils (introduced in [7]) and implemented it in a set of on-scalp MEG recordings using a 7-channel, high-*T*_c_ SQUID-based system [8]. The method provided high accuracy estimates of the sensor positions and orientations with short averaging time (≤ 2 mm and < 3 degrees respectively with 1-second coil recordings). It enables continuous estimation of the positions of sensors with respect to a subject’s head (i.e., head localization) with good temporal resolution. Calibrating and jointly localizing the sensor array can furthermore improve the localization accuracy (< 1 mm and < 2.5 degrees respectively with 1-second coil recordings). We demonstrate the efficacy of the method by using it in localization of neural activity.

## Acknowledgments

Data for this study was collected at NatMEG, the National infrastructure for Magnetoencephalography, Karolinska Institutet, Sweden. The NatMEG facility is supported by the Knut & Alice Wallenberg foundation (2011-0207). This work was financially supported by the Knut and Alice Wallenberg foundation (KAW 2014.0102), the Swedish Research Council (2017-00680), the Swedish Childhood Cancer Foundation (MT2014-0007), and Tillväxtverket via the European Regional Development Fund (20201637).

## References

[1] E. Boto, R. Bowtell, P. Krüger, T. M. Fromhold, P. G. Morris, S. S. Meyer, G. R. Barnes, M. J. Brookes, On the potential of a new generation of magnetometers for MEG: a beamformer simulation study, PLOS ONE 11 (8) (2016) e0157655.

[2] J. Iivanainen, M. Stenroos, L. Parkkonen, Measuring MEG closer to the brain: Performance of on-scalp sensor arrays, NeuroImage 147 (2017) 542–553.

[3] J. F. Schneiderman, S. Ruffieux, C. Pfeiffer, B. Riaz, On-scalp meg. in: Supek s., aine c. (eds) magnetoencephalography, Springer (2019) 1–23.

[4] B. Riaz, C. Pfeiffer, J. F. Schneiderman, Evaluation of realistic layouts for next generation on-scalp MEG: spatial information density maps, Scientific Reports 7 (1) (2017) 6974.

[5] S. Erné, L. Narici, V. Pizzella, G. Romani, The positioning problem in biomagnetic measurements: A solution for arrays of superconducting sensors, IEEE Transactions on Magnetics 23 (2) (1987) 1319–1322.

[6] K. Uutela, S. Taulu, M. Hämäläinen, Detecting and correcting for head movements in neuromagnetic measurements, NeuroImage 14 (6) (2001) 1424–1431.

[7] C. Pfeiffer, L. M. Andersen, D. Lundqvist, M. Hämäläinen, J. F. Schneiderman, R. Oostenveld, Localizing on-scalp MEG sensors using an array of magnetic dipole coils, PLOS ONE 13 (5) (2018) e0191111.

[8] C. Pfeiffer, S. Ruffieux, L. Jönsson, M. L. Chukharkin, A. Kalaboukhov, M. Xie, D. Winkler, J. F. Schneiderman, A 7-channel high-Tc SQUID-based on-scalp MEG system, bioRxiv (2019) 534107.

[9] M. Xie, J. F. Schneiderman, M. L. Chukharkin, A. Kalabukhov, B. Riaz, D. Lundqvist, S. Whitmarsh, M. Hämäläinen, V. Jousmäki, R. Oostenveld, et al., Benchmarking for on-scalp MEG sensors, IEEE Transactions on Biomedical Engineering 64 (6) (2017) 1270–1276.

[10] L. M. Andersen, R. Oostenveld, C. Pfeiffer, S. Ruffieux, V. Jousmäki, M. Hämäläinen, J. F. Schneiderman, D. Lundqvist, Similarities and differences between on-scalp and conventional in-helmet magnetoencephalography recordings, PLOS ONE 12 (7) (2017) e0178602.

[11] R. Oostenveld, P. Fries, E. Maris, J.-M. Schoffelen, Fieldtrip: Open source software for advanced analysis of MEG, EEG, and invasive electrophysiological data, Computational Intelligence and Neuroscience 2011 (2011) 1–9.

[12] R. Leahy, J. Mosher, M. Spencer, M. Huang, J. Lewine, A study of dipole localization accuracy for MEG and EEG using a human skull phantom, Electroencephalography and clinical neurophysiology 107 (2) (1998) 159–173.

[13] T. Bardouille, L. Power, M. Lalancette, R. Bishop, S. Beyea, M. J. Taylor, B. T. Dunkley, Variability and bias between magnetoencephalography systems in non-invasive localization of the primary somatosensory cortex, Clinical neurology and neurosurgery 171 (2018) 63–69.

[14] R. Zetter, J. Iivanainen, M. Stenroos, L. Parkkonen, Requirements for coregistration accuracy in on-scalp MEG, Brain topography 31 (6) (2018) 931–948.

[15] O. Alem, R. Mhaskar, R. Jiménez-Martίnez, D. Sheng, J. LeBlanc, L. Trahms, T. Sander, J. Kitching, S. Knappe, Magnetic field imaging with microfabricated optically-pumped magnetometers, Optics Express 25 (7) (2017) 7849–7858.

[16] J. Iivanainen, R. Zetter, M. Grön, K. Hakkarainen, L. Parkkonen, Onscalp MEG system utilizing an actively shielded array of optically-pumped magnetometers, NeuroImage 194 (2019) 244 –258.

[17] E. Boto, N. Holmes, J. Leggett, G. Roberts, V. Shah, S. S. Meyer, L. D. Muñoz, K. J. Mullinger, T. M. Tierney, S. Bestmann, et al., Moving magnetoencephalography towards real-world applications with a wearable system, Nature 555 (7698) (2018) 657.

[18] A. Borna, T. R. Carter, J. D. Goldberg, A. P. Colombo, Y.-Y. Jau, C. Berry, J. McKay, J. Stephen, M. Weisend, P. D. Schwindt, A 20-channel magnetoencephalography system based on optically pumped magnetometers, Physics in Medicine & Biology 62 (23) (2017) 8909.

